# Continuous attractors for dynamic memories

**DOI:** 10.1101/2020.11.08.373084

**Authors:** Davide Spalla, Isabel M. Cornacchia, Alessandro Treves

## Abstract

Episodic memory has a dynamic nature: when we recall past episodes, we retrieve not only their content, but also their temporal structure. The phenomenon of replay, in the hippocampus of mammals, offers a remarkable example of this temporal dynamics. However, most quantitative models of memory treat memories as static configurations, neglecting the temporal unfolding of the retrieval process. Here we introduce a continuous attractor network model with a memory-dependent asymmetric component in the synaptic connectivity, that spontaneously breaks the equilibrium of the memory configurations and produces dynamic retrieval. The detailed analysis of the model with analytical calculations and numerical simulations shows that it can robustly retrieve multiple dynamical memories, and that this feature is largely independent on the details of its implementation. By calculating the storage capacity we show that the dynamic component does not impair memory capacity, and can even enhance it in certain regimes.

## Introduction

The temporal unfolding of an event is an essential component of episodic memory. When we recall past events, or we imagine future ones, we do not produce static images but temporally structured movies, a phenomenon that has been referred to as “mental time travel” [1], [2].

The study of the neural activity of the hippocampus, known for its first-hand involvement in episodic memory, has provided many insights on the neural basis of memory retrieval and its temporal dynamics. An interesting example is the phenomenon of hippocampal replay, i.e. the reactivation, on a compressed time scale, of sequences of cells active in previous behavioural sessions. Replay takes place during sharp wave ripples, fast oscillations of the hippocampal local field potential that are particularly abundant during sleep and restful wakefulness [3], [4]. Indeed, replay has been observed during sleep [5], [6], inter-trial rest periods [7], [8], and during still periods in navigational tasks [9], [10]. Replay activity has been hypothesized to be crucial for memory consolidation [11] and retrieval [12], as well as for route planning [10], [13].

A temporally structured activation takes place also before the exposure to an environment [14], a phenomenon known as *preplay*, and a recent study showed that this dynamical feature emerges very early during development, preceding the appearance of theta rhythm [15] in the hippocampus. The fact that hippocampal sequences are present before the exposure to the environment they will encode suggests that their dynamical nature is not specific to a role in spatial cognition, but is inherent to hippocampal operation in general. Moreover, in a recent study Stella et al. [16] have shown that the retrieved sequences of positions during slow wave sleep are not always replaying experienced trajectories, but are compatible with a random walk on the low dimensional manifold that represents the previously explored environment. This suggests that what is essential are not the sequences themselves, but the tendency to produce them: neural activity tends to move, constrained to abstract low-dimensional manifolds, which can then be recycled to represent spatial environments, and possibly non-spatial ones as well. This dynamic nature extends on multiple timescales, as suggested by the observation that the neural map of the same environment progressively changes his components cells over time [17].

Low-dimensional, dynamic activity is not constrained to a single subspace: replay in sleep can reflect multiple environments [18], [19], the content of awake replay reflects both the current and previous environments [12], and during behaviour fast hippocampal sequences appear to switch between possible future trajectories [20]. Further evidence comes from a recent study with human participants learning novel word pair associations [21]. The study shows that the same, pair-dependent neural sequences are played during the encoding and the retrieval phase.

A similar phenomenon – a dynamic activity on low dimensional manifolds – is present in memory schemata, cognitive frameworks that constrain and organize our mental activity [22], and have been shown to have a representation in the medial temporal lobe [23]. Yet another example of dynamical, continuous memories is offered by motor programs, which have been described as low-dimensional, temporally structured neural trajectories [24], [25], [26], or as dynamical flows on manifolds [27], [28].

We refer to these objects as *dynamical continuous attractors*, since they involve a continuous subspace that constrains and attracts the neural activity, and a dynamical evolution in this subspace. Fig.1 schematically illustrates the concept of dynamical continuous attractors and their possible role in some of the neural processes described above.

**Figure 1:**
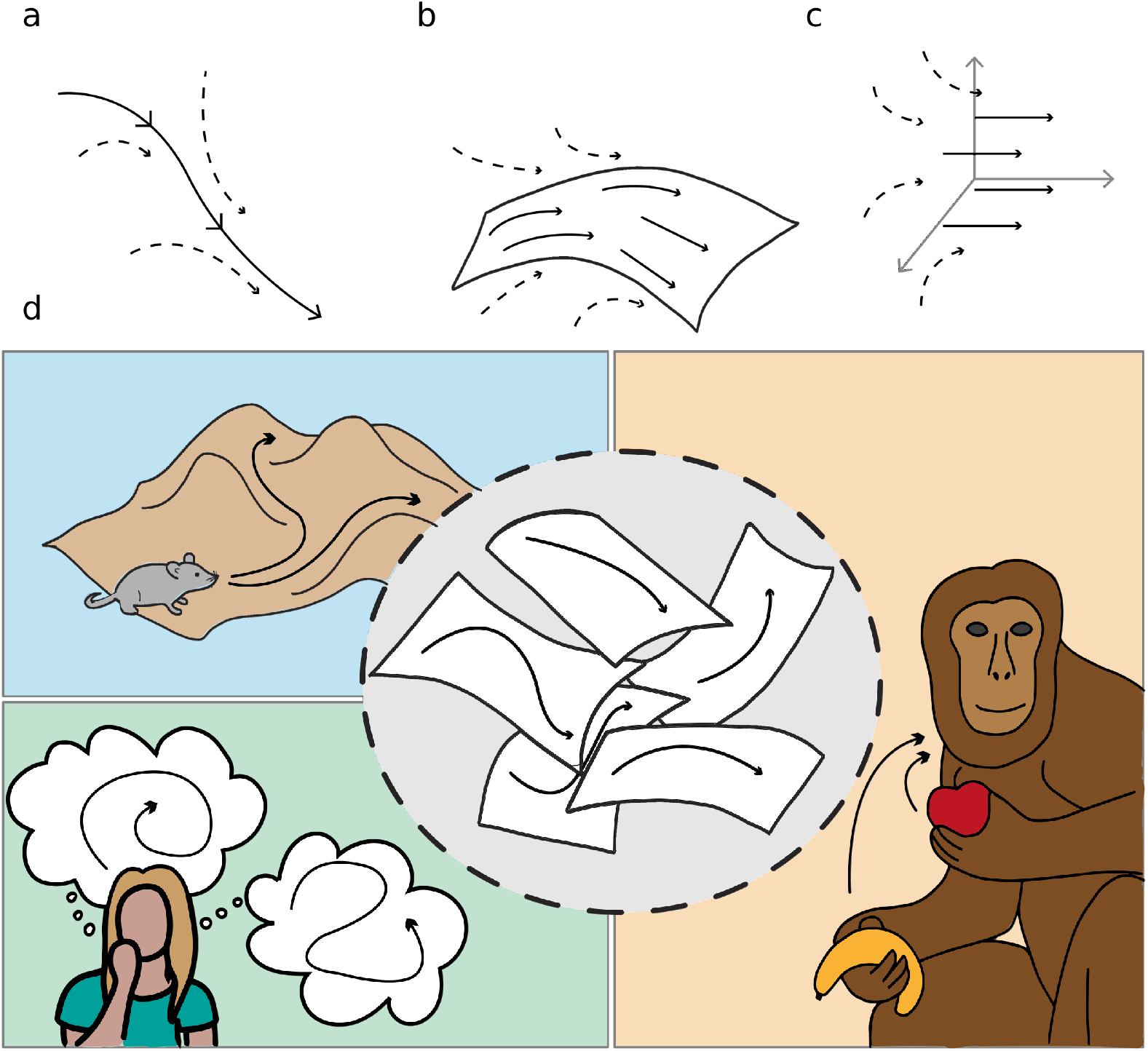
Schematic illustration of dynamic continuous attractors as a basis of different neural processes. Top row: a scheme of continuous attractive manifolds, with a dynamic component in 1D **(a)**, 2D **(b)** and 3D **(c)**. The neural activity quickly converges on the attractive manifold (dotted arrows), then slides along it (full arrows), producing a dynamics that is temporally structured and constrained to a low dimensional subspace. Bottom row: multiple dynamic memories could be useful for route planning (top left), involved in mind wandering activity (bottom left) or represent multiple learned motor programs (right).

In most cases, computational analyses of low-dimensional neural dynamics are not concerned with memory, and focus on the description of the features of single attractors, more than on their possible coexistence. On the other hand, mechanistic models of memory usually neglect dynamical aspects, treating memories as static objects, either discrete [29] or continuous [30], [31], [32], [33]. The production of sequences of discrete memories can be implemented with a heteroassociative component [34], usually dependent on the time integral of the instantaneous activity, that brings the network out of equilibrium and to the next step in the sequence. A similar effect can be obtained with an adaptation mechanism in a coarse grained model of cortical networks [35], with the difference that in this case the transitions are not imposed, but driven by the correlations between the memories in so-called latching dynamics [36], [37]. Moreover, adaptation-based mechanisms have been used to model the production of random sequences on continuous manifolds [38], and shown to be crucial in determining the balance between retrieval and prediction in a network describing CA3-CA1 interactions [39]. In the case of continuous attractor networks, movement can be induced also by mechanisms that integrate an external velocity input and make use of asymmetric synaptic strengths. Models of this kind have been used for the description of head direction cells [40], spatial view cells [41], and grid cells [42], [43], and can represent simultaneously the positions of multiple features and their temporal evolution [44]. In the simplest instantiation, these systems do not necessarily reflect long-term memory storage: the activity is constrained on a single attractive manifold, which could well be experience independent.

Here we propose a network model able to store and retrieve multiple, independent dynamic continous attractors. The model relies on a map-dependent asymmetric component in the connectivity, that produces a robust shift of the activity on the retrieved attractive manifold. This connectivity profile is conceived to be the result of a learning phase in which the mechanism of spike timing dependent plasticity (STDP) [45] produces the asymmetry. Crucially, the asymmetry is not treated here as a “pathological” feature, assumed to level out in the limit of long learning, but as a defining trait of the stored memories. The balance between two components – one symmetric and trajectory-averaged, the other asymmetric and trajectory-dependent – is explicit in the formulation of the model, and allows to study their effects on memory storage.

In what follows we develop an analytical framework that allows to derive the dependence of important features of the dynamics, such as the replay speed and the asymmetry of the activity cluster, as a function of the relevant parameters of the model. We show with numerical simulations that the behaviour of the model is robust with respect to its details, and depends weakly on the shape of the interactions. Finally, we estimate the storage capacity for dynamical memories and we find it to be of the same order of the capacity for static continuous attractors, and even higher in some regimes.

### A mechanistic model for dynamic retrieval

The model we consider is a continuous attractor neural network, with an additional anti-symmetric component in the connectivity strength. We consider a population of *N* neurons, with recurrent connectivity described by an interaction matrix *J*_*ij*_, whose entries represent the strength of the interaction between neuron *i* and *j*. The activity of each neuron is described by a positive real number *V*_*i*_∈ ℝ^+^representing its instantaneous firing rate. The dynamic evolution of the network is regulated by the equations:

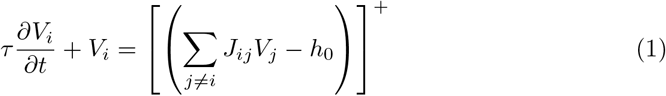

where […]^+^ is the threshold linear activation function

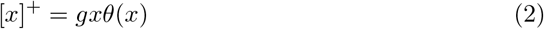

with the gain *g* modulating the slope and the Heaviside step function *θ*(…) setting to zero sub-threshold inputs. The first term on the right hand side of eq. 1 represent the excitatory inputs provided to neuron *i* from the rest of the network through recurrent connections. The threshold *h*_0_ and the gain *g* are global parameters that regulate the average activity and the sparsity of the activity pattern [46].

In numerical simulations, these parameters are dynamically adjusted at each time step to constrain the network to operate at a certain average activity (usually fixed to 1 without loss of generality) and at a certain sparsity *f*, defined as the fraction of active neuron at each time (see appendix A). The connectivity matrix *J* of the network encodes a map of a continuous parameter 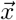 spanning a low-dimensional manifold, e.g. the position in an environment. To do so, each neuron is assigned a preferential firing location 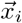 in the manifold to encode, and the strength of the interaction between pairs of neuron is a given by a decreasing, symmetric function of the distance between their preferred firing locations

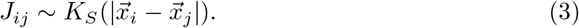

This shape of the interactions is a typical one in the framework of continuous attractor neural networks [32] [30] [47], and is thought to come from a time-averaged Hebbian plasticity rule: neurons with nearby firing fields will fire concurrently and strengthen their connections, while firing fields far apart will produce weak interactions. The symmetry of the function *K*_*S*_, usually called **interaction kernel**, ensures that the network reaches a static equilibrium, where the activity of the neurons represents a certain position in the map and, if not pushed, remains still.

### The shift mechanism

The assumption of symmetric interactions neglects any temporal structure in the learning phase. In case of learning a spatial map, for example, the order in which recruited neurons fire along a trajectory may produce an asymmetry in the interactions as a consequence of Spike Timing Dependent Plasticity [45], that requires the postsynaptic neuron to fire *after* the presynaptic one in order to strengthen the synapse. This phenomenon can be accounted for in the definition of the interaction kernel. Any asymmetric kernel can be decomposed in two contributions:

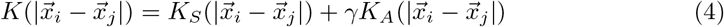

where *K*_*S*_ is the usual symmetric component and *K*_*A*_ is an anti-symmetric function (*K*_*A*_(*x*_*i*_ − *x*_*j*_) = −*K*_*A*_(*x*_*j*_ − *x*_*i*_)). The parameter *γ* regulates the relative strength of the two components. The presence of *K*_*A*_ will generate a flow of activity along the direction of asymmetry: neuron *i* activates neuron *j* that, instead of reciprocating, will activate neurons downstream in the asymmetric direction. Mechanisms of this kind have been shown to produce a rigid shift of the encoded position along the manifold, without loss of coherence [40], [43], [42]. In the quantitative analysis that follows we will concentrate, when not stated otherwise, on a kernel *K* with the exponential form

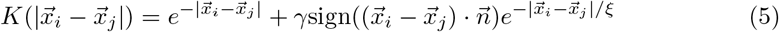

Where 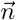 is a unit vector pointing in the – constant – direction along which the asymmetry is enforced, and *ξ* is the spatial scale of the asymmetric component, fixed to 1 where not explicitly stated otherwise. This form simplifies the analytical description of the model, but is not special in any way. In fact, all the result presented hold for a large classes of interaction kernels. The robustness of the model with respect to the details of the kernel is specifically addressed below.

### Asymmetric recurrent connections produce dynamic retrieval

The spontaneous dynamics produced by the network is constrained on the low dimensional manifold codified in the connectivity matrix and spanned by the parameter 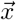. The short-range interactions and the uniform inhibition enforced by the firing threshold *h*_0_ produce a localized “bump” of activity in the manifold. The presence of an asymmetry in the connection strengths prevents the system from remaining in a stationary equilibrium. Instead, it generates a steady flow of activity in the direction of the asymmetry. This flow is illustrated in Fig. 2 (a),(b) and (c), obtained with numerical simulation of a network encoding a one-, two- or three-dimensional manifolds, respectively: the activity clusters in a bump around the encoded position at each time point (*t*_1_,*t*_2_ and *t*_3_), and the bump shifts without dissipation by effect of the asymmetric component of the interactions. The coherence of the representation is not affected by the dynamics: the network is able to retrieve a dynamically evolving position on the continuous manifold spanned by 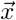.

**Figure 2:**
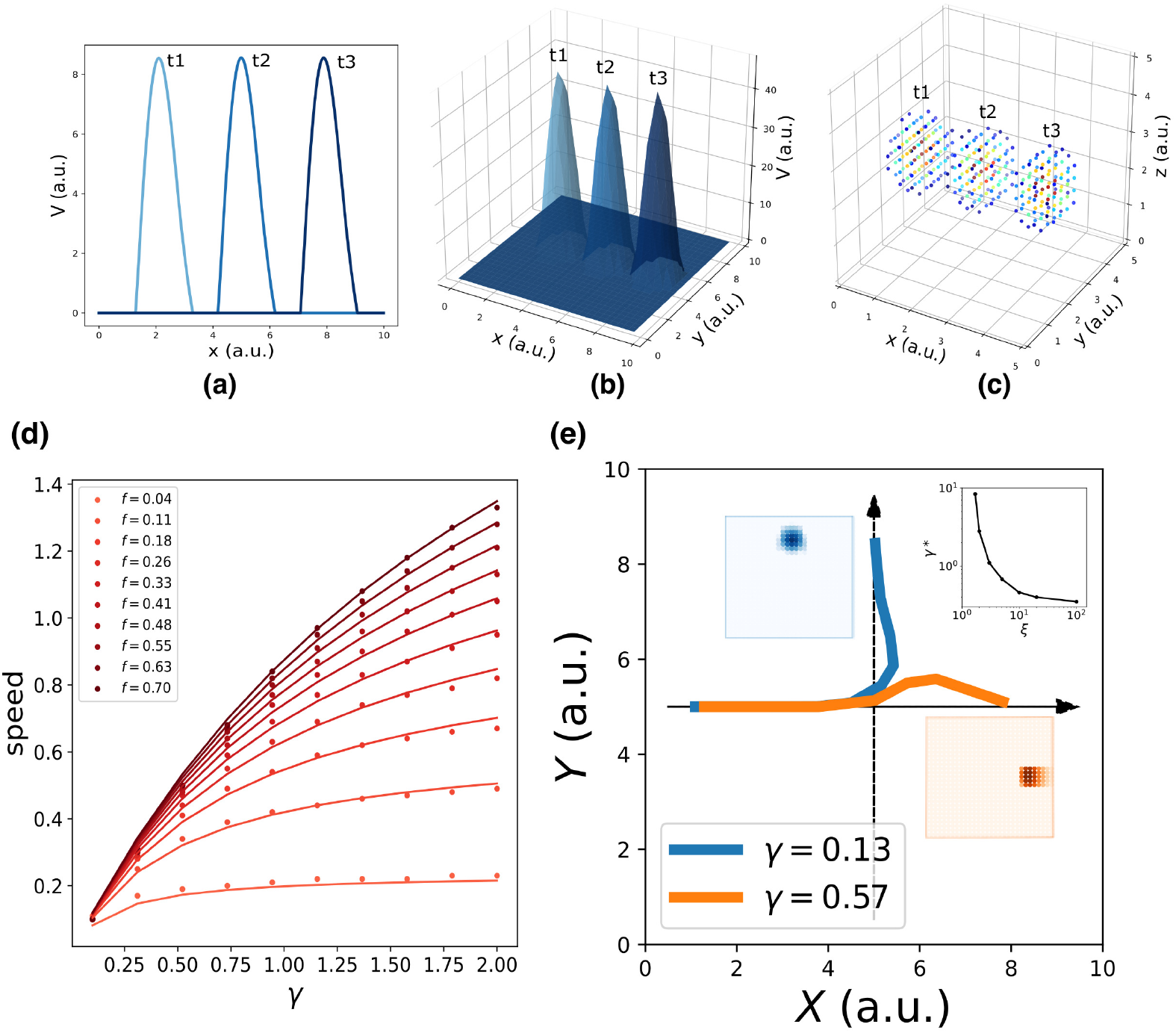
Dynamic retrieval of a continuous manifold. First row: in each plot are three snapshots of the network activity at three different times (t1, t2 and t3), for a system encoding a one dimensional **(a)**, two dimensional **(b)** and three dimensional manifold **(c)**. In (c), activity is color-coded (blue represents low activity, red high activity, silent neurons are not plotted for better readability). In all cases the anti-symmetric component is oriented along the x axis. **(d)** Dependence of the speed on *γ* and *f*. Dots are data from numerical simulations, full lines the fitted curves. **(e)** Retrieval of two crossing trajectories: full line show the trajectories followed by the center of mass of the activity. Blue curve: low *γ*, switch between trajectories; Orange curve: high *γ*, successful crossing. In both cases *ξ* = 10. The blue and the orange insets show the activity bumps in the corresponding cases; the top-right inset shows the dependence of the value *γ*^*\**^, required for crossing, on *ξ*.

The speed of movement of the bump is modulated by the value of the parameter *γ* and by the sparsity level of the representation *f*, i.e. the fraction of neurons active at each given time during the dynamics. The dependence of the speed on these two parameters is illustrated in 2(d). Stronger asymmetry (high *γ*) produce a faster shift. Interestingly, the sparsity value *f* acts as a modulator of the influence of *γ*: sparser representations move more slowly than dense ones.

While *γ* describes a feature of the synaptic interactions, determined during the learning phase and relatively fixed at the short timescales of retrieval, *f* can be instantaneously modulated during retrieval dynamics. A change in the gain or the excitability of the population can be used to produce dynamic retrieval at different speeds. Thus, the model predicts an interaction between the sparsity and the speed of the reactivation of a continuous memory sequence, with increased activity leading to faster replay. It is worth noting, however, that the *direction* of the dynamics is fixed with *J* : the model is able to retrieve either forward or backward sequences, but not alternate between them.

The dependence of the retrieval speed *s* on *γ* and *f* is well described by the approximate functional form

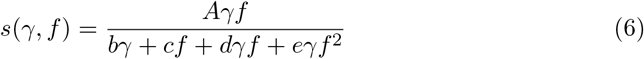

This dependence is shown in 2(b), where the dots are the values obtained with numerical simulations and the full curves the fitted relationship. The full understanding of the nature of this functional form remains an open challenge for future analysis. As we will see in the next section, the analytical solution of the model yields the same form to describe the dependence of speed on *γ* and *f*, but a closed-form solution is still lacking.

In the model presented, the asymmetry in the interactions is enforced uniformly along a single direction also for two- and three-dimensional manifolds, representing the case in which neural dynamics follows a forced trajectory along one dimension, but is free to move without energy costs along the other. However, the same mechanism can be used to produce one-dimensional trajectories embedded in low-dimensional manifolds, with the introduction of a positional dependence in the direction of the asymmetry [48]. In this case, an interesting problem is posed by the intersection of two trajectory embedded in the same manifold: is the network, during the retrieval of one trajectory, able to cross these intersections, or do they hinder dynamical retrieval? We present in Fig. 2 (e) a numerical study of a minimal version of this problem, with two orthogonal trajectories (black arrows) embedded in a 2D manifold and memorized simultaneously in the network. When the network is cued to retrieve the horizontal trajectory, the behaviour at the intersection depends on the strength *γ* and scale *ξ* of the asymmetric component. At low *γ* the dynamics spontaneously switch trajectory at the intersection (Fig 2(e), blue curve), while for *γ* sufficiently large the retrieval of the horizontal trajectory is successful (Fig, 2(e), orange curve). The value *γ*\* required for a successful crossing depends on the spatial scale *ξ*: larger *ξ* allow for crossing with lower values of *γ*, as shown in the top-right inset of Fig. 2(e), in which *γ*\* (*ξ*) is plotted. Intuitively, the ability of the network to retrieve crossing trajectories depends on the shape of the activity bump, that needs to be sufficiently elongated in the direction of retrieval for the succesful crossing of the intersection. The blue and orange insets of Fig. 2(e) show the difference in shape of the bump in the case of a trajectory switch (blue) and crossing (orange).

### Analytical solution for the single manifold case

The simplicity of continuous attractor models often allows to extract important computational principles from their analytical solution [49] [50]. In our case, the dynamic behaviour of the system and its features can be fully described analytically with a generalization of the framework developed by Battaglia & Treves [30]. For this purpose, it is easier to formulate the problem in the continuum, and describe the population activity {*V*_*i*_} by its profile 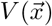 on the attractive manifold parametrized by the coordinate 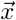, and the dynamical evolution as a discrete step map, equivalent to Eq. 1.

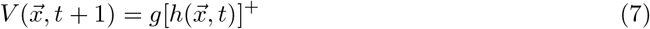

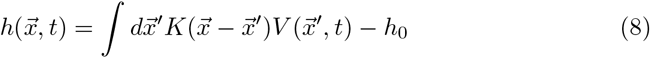

The requirement of a rigid shift of population activity is then imposed by setting the activity at time *t* + 1 to be equal at the activity at time *t*, but translated by an amount 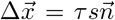, proportional to the speed *s* of the shift and in the direction 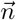 of the asymmetry in the connections. The timescale *τ* sets the time unit in which the duration of the evolution is measured and does not have an impact on the behaviour of the system.

The activity profile 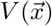 is then found as the self-consistent solution to the integral equation

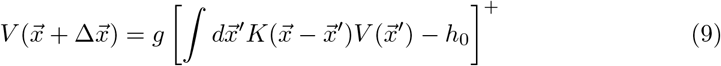

Equation 9 is valid in general. We will focus here, for the explicit derivation (reported in appendix B), on the case of a one dimensional manifold with an exponential interaction kernel

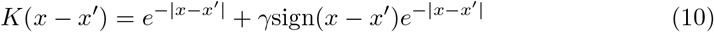

In this case, the activity bump will take the form:

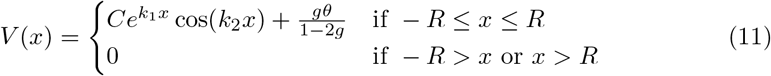

The parameters *k*_1_ = *k*_1_(*γ, s*) and *k*_2_ = *k*_2_(*γ, s*) determine the properties of the solution and depend on the value of *γ* and speed *s* = Δ*x/τ*.

*k*_2_ is related to the bump width by the relation

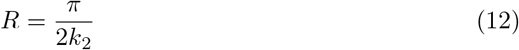

where *R* is the point at which *V* (*x*) = 0. *k*_1_ is related to the asymmetry of the bump: in the limit case *γ* = 0, *s* = 0 (Fig. 3(a), first column) *k*_1_ = 0, and we recover the cosine solution of the symmetric kernel case studied in [30]. Larger *k*_1_ values result in more and more asymmetric shapes (Fig. 3(a), second and third columns).

**Figure 3:**
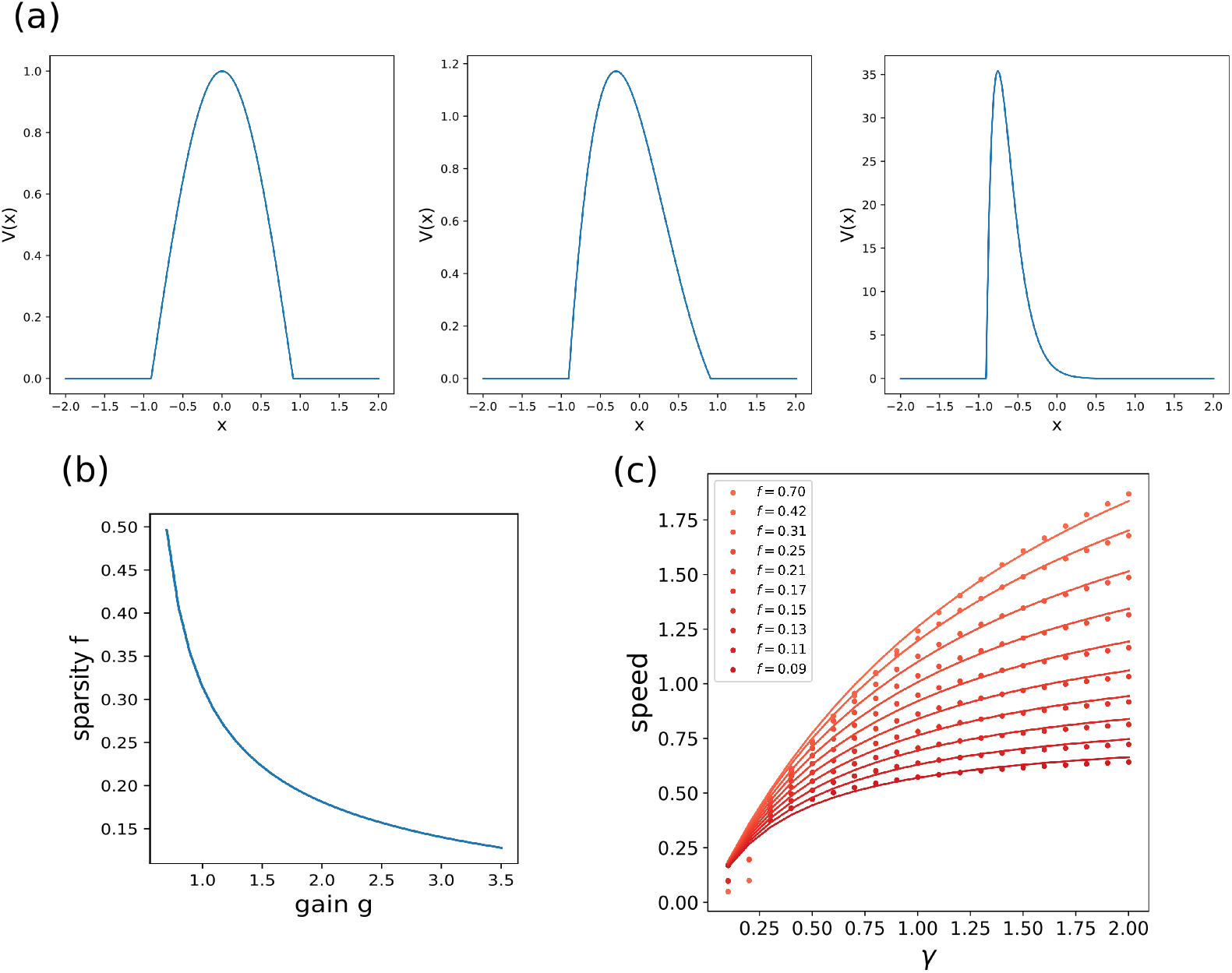
Analytical solution of the model. **(a)** The shape of the bump for increasing values of *γ*. **(b)** dependence of the sparsity *f* on the gain *g* of the network. **(c)** Dependence of the speed of the shift on *γ*, at different values of sparsity. Dots show the numerical solution (note some numerical instability at low *f* and *γ*), full curves the best fits.

From this analytical solution we can determine the dependence of the speed *s* on the asymmetry strength *γ* and on the sparsity *f* = 2*R/L* (note the in the continuum case the fraction of active neurons is given by the ratio between the bump size 2*R* and the manifold size *L*). The sparsity *f* is modulated by the value of the gain *g*, as shown in figure 3(b): a larger gain in the transfer function corresponds to a sparser activity. The exact relation *s*(*γ, f*) can be obtained from the numerical solution of a trascendental equation (see appendix B), and can be approximated with a functional shape analogous to the one used for the simulated network:

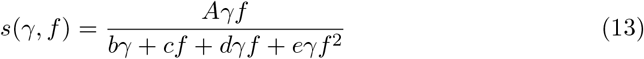

The full trascendental solution and the fitted curves are reported in Fig. 3(c).

### Dynamic retrieval is robust

The full analytical description of the model is presented here for a specific kernel choice, but the dynamic retrieval the network shows is extremely robust with respect to the details of the model implementation. Performing numerical simulations with different kernel choices, we find that a coherently shifting bump can be obtained with a wide variety of interaction kernels, without any relationship required, for example, between the shapes of the symmetric and anti-symmetric components. Some examples are illustrated in Fig. 4.

**Figure 4:**
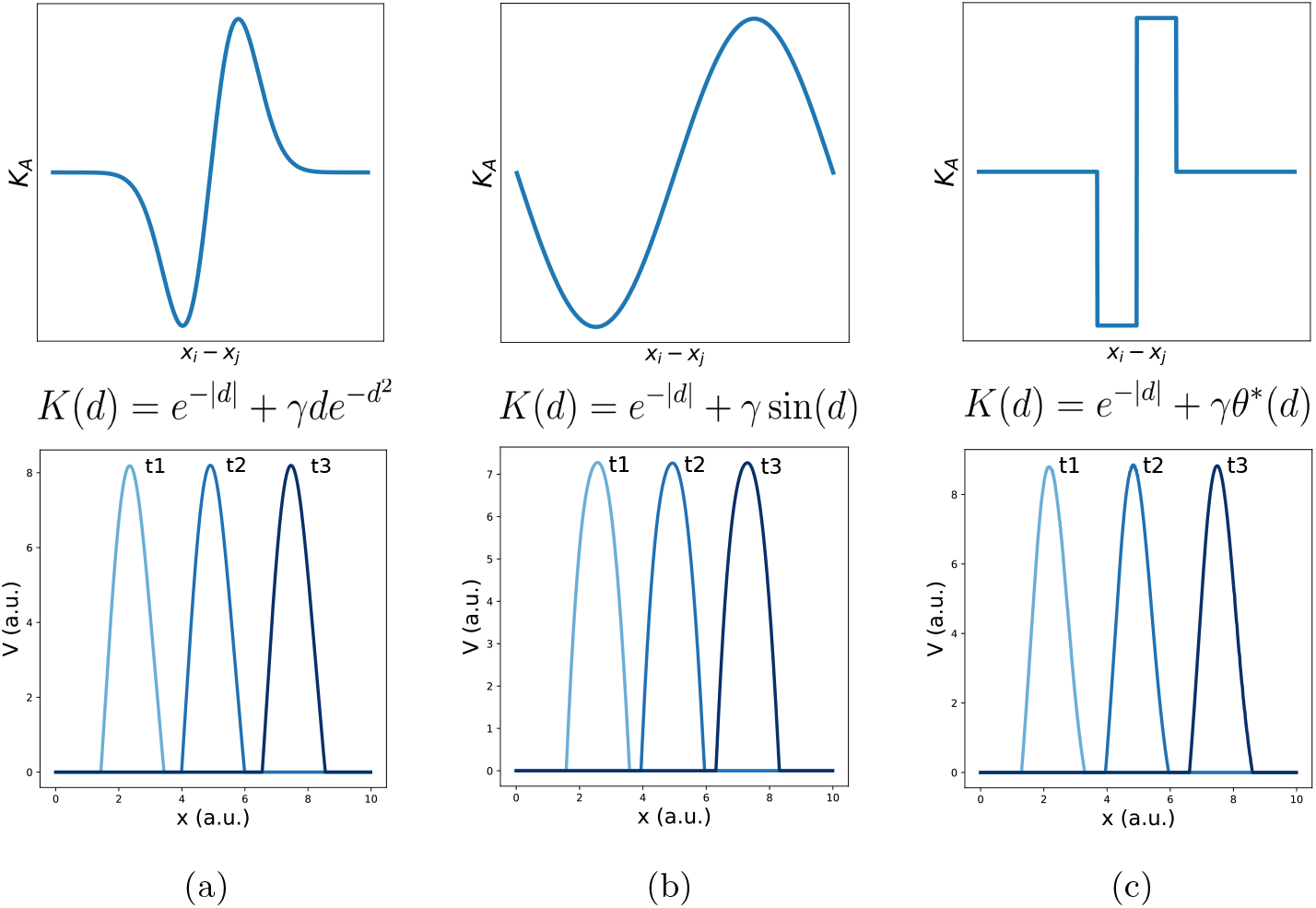
Different interaction kernels produce similar behaviour. Three examples of dynamics with the same symmetric component and three different anti-symmetric components. Top row: shape of the anti-symmetric component *K*_*A*_. Bottom row: three snapshots of the retrieval dynamics for the corresponding *K*_*A*_. **(a)** Gaussian derivative; Sinusoidal; **(c)** Anti-symmetric step function, *θ*^*\**^ = *θ*(*d*)*θ*(1 − *d*) − *θ*(−*d*)*θ*(*d* − 1).

We find two parameters to be important in determining the dynamical behaviour of the model: the relative strength *γ* between the symmetric and anti-symmetric components, and the characteristic length *ξ* of the anti-symmetric component. Their effect on the dynamics are shown in Fig. 5, obtained with numerical simulation of the network with different values of *γ* and *ξ*. In the whole the range of parameters analyzed, spanning four orders of magnitude in both parameters, the network was able to produce dynamic retrieval. *γ* and *ξ* affect the speed of the shift, the peak values of the activity distribution and the skewness of the activity bump, without hindering network functionality. Dynamical retrieval does not require any fine tuning of network parameters, nor a specific functional shape of the interaction kernel.

**Figure 5:**
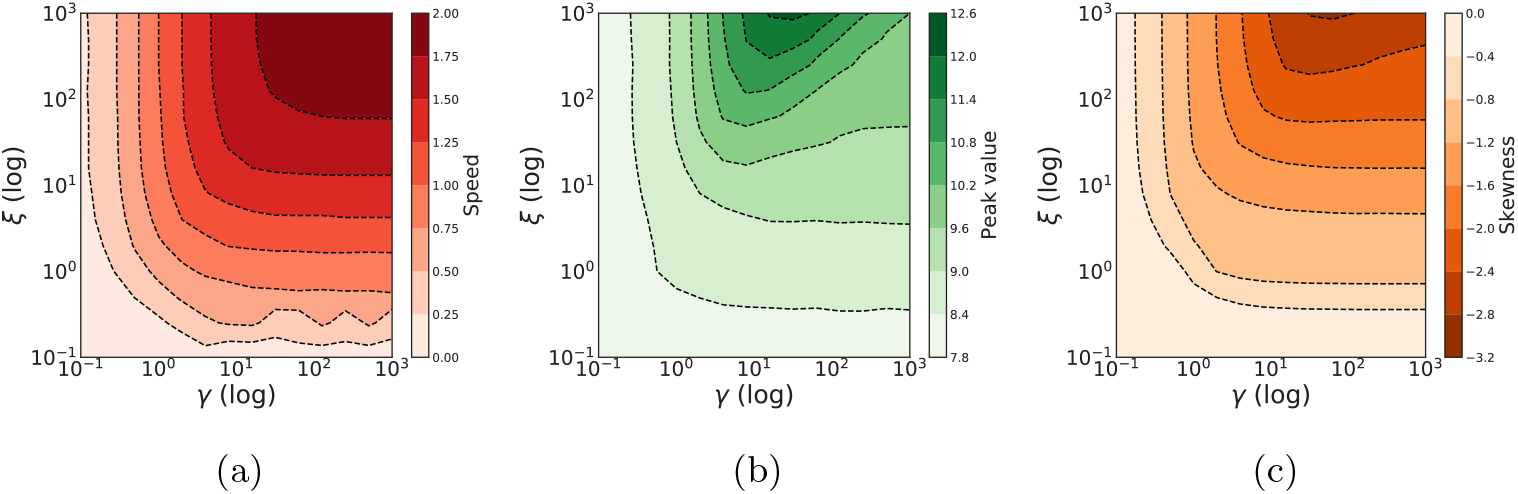
Dynamical retrieval in a wide range of parameters. Effect of the kernel strength *γ* and its spatial scale *ξ*, in the case of the exponential kernel *K*(*d*) = *e*^−|*d*|^ + *γsign*(*d*)*e*^−|*d*|*/ξ*^ .**(a)** Retrieval speed **(e)** Peak value of the activity **(e)** Skewness of the activity bump.

### A dynamical memory: storing multiple manifolds

We described in detail the behaviour of a neural network with asymmetric connectivity in the case of a single manifold encoded in the synaptic connectivity. For the network to behave as an autoassociative memory, however, it needs to be able to store and dynamically retrieve *multiple* manifolds. This is possible if we construct the interaction matrix *J*_*ij*_ as the sum of the contributions from *p* different, independently encoded manifolds:

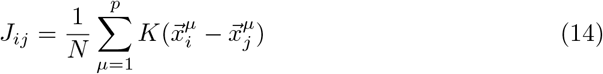

Here each 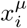 represent the preferred firing location of neuron *i* in the manifold *µ*, and *K* is the same interaction kernel as in eq. 4, containing a symmetric and anti-symmetric component.

The resulting dynamics shows multiple continuous attractors, corresponding to the stored manifolds. Given an initial configuration, the networks rapidly converges to the nearest (i.e. most correlated) attractor, forming a coherent bump that then moves along the manifold as a consequence of the asymmetric component of the connectivity. The same dynamics, if projected on the other, unretrieved manifolds, appears as random noise. This is illustrated in Fig.6 obtained with numerical simulations of a network encoding three different manifolds (of dimension 1 in (a), dimension 2 in (b)), and dynamically retrieving the first one.

**Figure 6:**
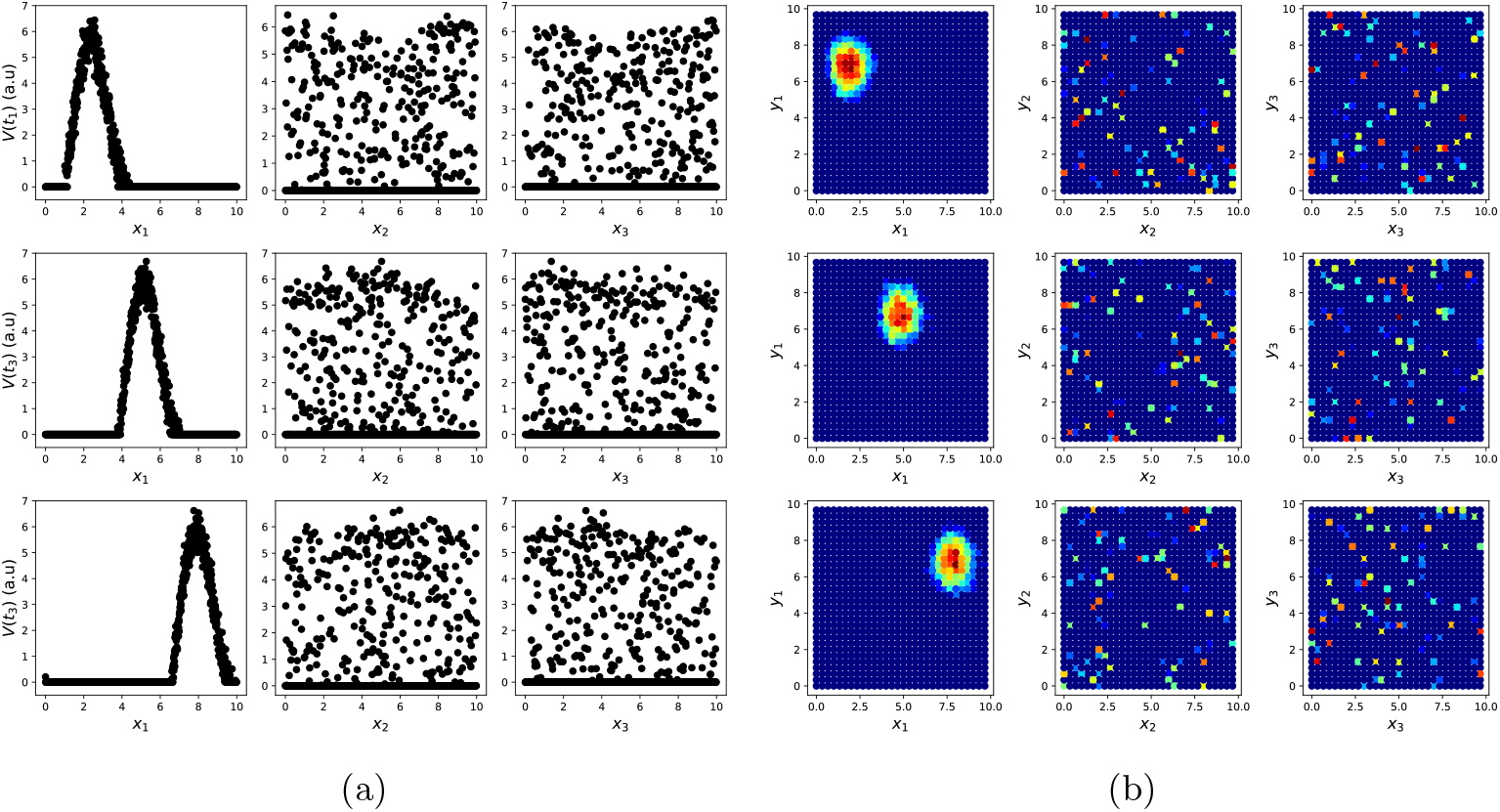
Dynamic retrieval in the presence of multiple memories. **(a)** In one dimension **(b)** In two dimensions. Each row represents a snapshot of the dynamics at a point in time. The activity is projected on each of the three attractors stored in the network. In both cases, the first attractor is retrieved, and the activity organizes in a coherent bump that shifts in time. The same activity, projected onto the two non-retrieved maps looks like incoherent noise ((a) and (b), second and third columns).

Multiple dynamic manifolds can be memorized and retrieved by the network, with different speeds. Fig. 7 (b) shows the result of the numerical simulation of a network with five different one-dimensional manifold stored in its connectivity matrix, each encoded with a different value of *γ* (see appendix C). These manifold are dynamically retrieved by the network at different speeds, depending on the corresponding *γ*. This allows the model to simultaneously store memories without the constraint of a fixed dynamical timescale, an important feature for the description of biological circuits that need to be able to operate at different temporal scales.

**Figure 7:**
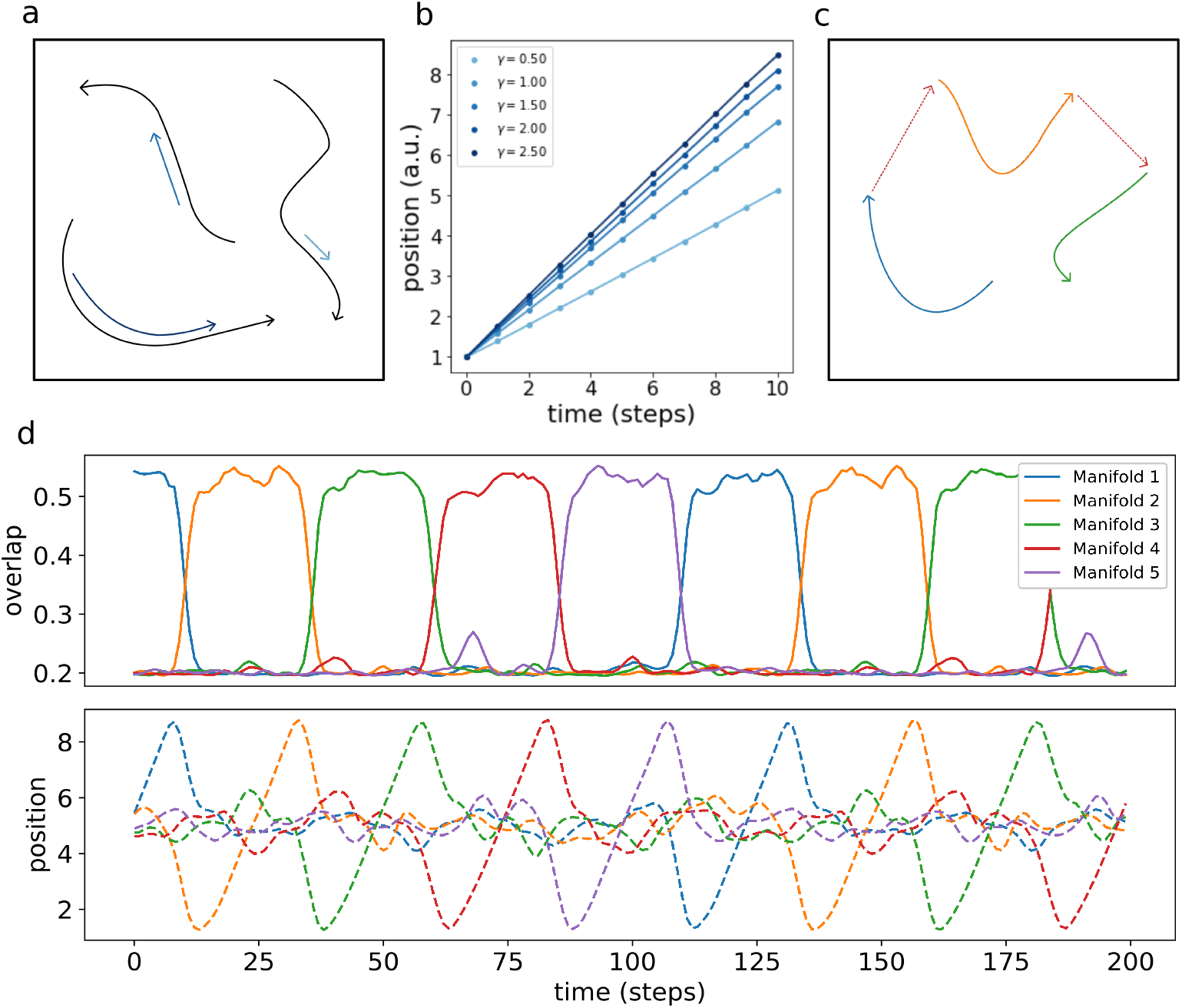
Retrieval speed and memory interactions. **(a)** Multiple mainfolds with different velocity can be stored in a network with manifold-dependent asymmetric connectivity **(b)** The retrieved position at different timesteps during the retrieval dynamics of five different manifolds, stored in the same network, each with a different value of *γ*. **(c)** Manifolds memorized in the same network can be linked together **(d)** Sequential retrieval of five manifolds. Top row: overlap, meausing the overall coherence with the manifold, as a function of time. Bottom row: retrieved position in each manifold as a function of time.

Different memories stored in the same neural population can interact with each other, building more retrieval schemes in which, for example, the retrieval of a memories cues the retrieval of another one. To investigate this possibility, we have incorporated in the model a mechanism for interaction between memories, in which the endpoint of a dynamical, one-dimensional manifold elicits the activation of the start point of a different one (see appendix D). This results in the sequential retrieval of multiple memories, one after the other, as illustrated in Fig. 7 (d). The top row shows the evolution in time of the overlaps *m*_*µ*_:

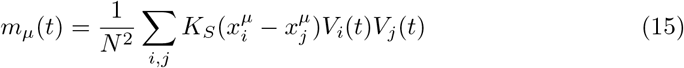

These order parameters quantify the coherence of the population activity *V* (*t*) with each of the manifolds. Localized activity in manifold *µ* results in a large *m*_*µ*_, while a low *m*_*µ*_ corresponds to an incoherent scattering of the activity. The network retrieves the manifolds in sequence, one at a time, following the instructed transitions encoded in its connectivity. The all-or-none behaviour of the coherence parameters segments the continuous dynamics of the network into a sequence of discrete states.

The bottom row shows the evolution of the retrieved position, given in each manifold by the center of mass:

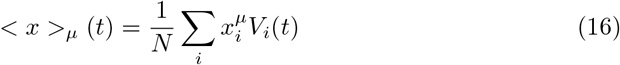

The dynamics runs across the retrieved manifold, from its beginning to its end, then jumps to next one and repeats the process. Note that the position in each of the unretrieved manifold fluctuates around *L/*2, as a consequence of the incoherence of the activity. Within each of the retrieved manifolds the dynamics retains its continuous nature in the representation of the evolving position.

This sequential dynamics goes beyond the simple cued retrieval of independent memories that is the focus of most autoassociative memory models, and provides an example of a hybrid computational system, encoding both continuous and discrete features.

The interaction mechanism introduced here provide the opportunity to investigate the effect of more complex interactions than the simple memory chain presented here. We present here this first example as a proof of principle of the possibility of storing interacting dynamical memories, and will proceed to the investigation of more complex structures (e.g. interaction networks, probabilistic interactions, etc..) in future studies.

### Storage capacity

The number of maps that can be stored and retrieved by an attractor network of this kind is typically proportional to the number of inputs per neuron *C* [51]. The memory load *α* = *p/C* crucially determines the behaviour of the system: when *α* is increased above a certain threshold value *α*_*c*_, the network is not able to retrieve any of the stored memories, falling instead into a disordered state. Therefore it is the magnitude of *α*_*c*_, i.e. the storage capacity of the system, that determines how effectively it can operate as a memory. To estimate the storage capacity for dynamic continuous attractors, and to investigate how it is impacted by the presence of asymmetric connections, we proceed along two complementary paths.

In the case a fully connected network, where the analytical tools developed for equilibrium systems are not applicable, we take advantage of the fact that numerical simulations can be effective for the estimation of the capacity, since the number of connections per neuron *C* (the relevant parameters in the definition of the storage capacity *α*_*c*_ = *p/C*) coincides with the number of neurons, minus one. For a highly diluted system, on the other hand, the number of neurons is much larger than *C*, making the simulation of the system very difficult in practice. We then resort to an analytical formulation based on a signal to noise analysis [30], that exploits the vanishing correlations between inputs of different neurons in a highly diluted network, and does not require symmetry in the connectivity [52]. The quantification of the effect of loops in the dense connectivity regime, developed in [53] and [54] for the case of static, discrete attractors, is beyond the scope of the present work and remains an interesting open direction.

In both the fully connected and the highly diluted case we study the dependence of the capacity on two important parameters: the map sparsity, i.e. the ratio between the width of the connectivity kernel (fixed to one without loss of generality) and the size *L* of the stored manifolds, and the asymmetry strength *γ*. Note that the map sparsity 1*/L* is different from the activity sparsity *f*: the former is a feature of the stored memories, that we will treat as a control parameter in the following analysis; the latter is a feature of the network dynamics, and its value will be fixed by an optimization procedure in the calculation of the maximal capacity.

### Analytical calculation of the capacity in the highly diluted limit

The signal-to-noise approach we follow, illustrated in details in [30], involves writing the local field *h*_*i*_ as the sum of two contributions: a signal term, due to the retrieved – “condensed” – map, and a noise term consisting of the sum of the contributions of all the other, “uncondensed” maps. In the diluted regime (*C*/*N* → 0) these contributions are independent and can be summarized by a Gaussian term *ρz*, where *z* is a random variable with zero mean and unit variance. In the continuous limit, assuming without loss of generality that map *µ* = 1 is retrieved, we can write:

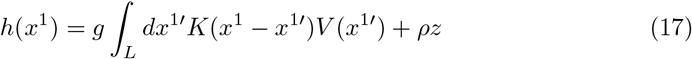

The noise will have variance:

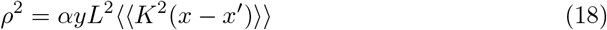

Where *L* is the size of the map, ⟨⟨*K*^2^(*x* − *x*′)⟩⟩ is the spatial variance of the kernel and

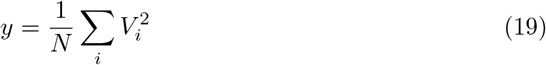

is the average square activity.

We can write the fixed point equation for the average activity profile *m*^1^(*x*), incorporating the dynamic shift with an argument similar to the one made for the single map case:

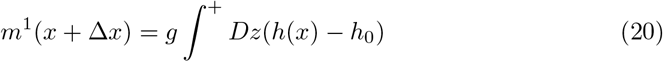

Where 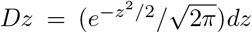 and ∫^+^ *f*(*x*)*dx* = ∫ *f*(*x*)*θ*(*x*)*dx*. The average square activity *y*, entering the noise term, reads

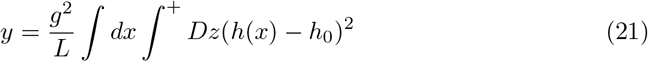

Introducing the rescaled variables

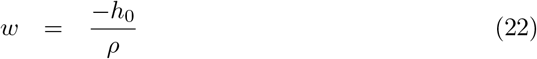

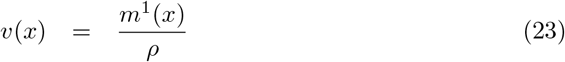

And the functions

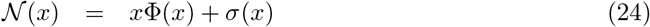

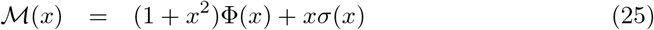

where Φ(*x*) and *σ*(*x*) are the Gaussian cumulative and the Gaussian probability mass function respectively, we can rewrite the fixed-point equation as

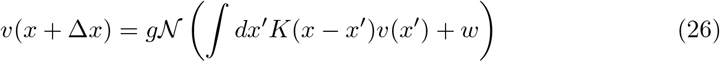

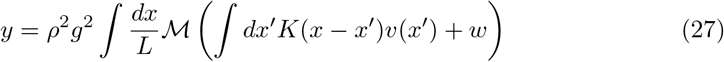

Substituting Eq.27 in the expression for the noise variance 18 we obtain

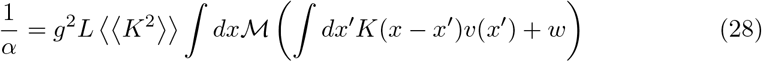

If we are able to solve Eq. 26 for the rescaled activity profile *v*(*x*), we can use Eq. 28 to calculate *α*. We can then maximize *α* with respect to *g* and *w*: this yields the maximal value *α*_*c*_ for which retrieval solutions can be found.

These equations are valid in general and have to be solved numerically. Here we present the results for the case of one-dimensional manifolds and interactions given by the exponential kernel of Eq. 35. In this case we have

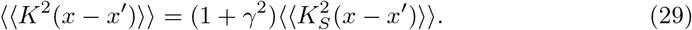

where *K*_*S*_(*x*−*x*′) = *e*^−|*x*−*x′* |^ is the symmetric component of the kernel. A simple approximation, illustrated in appendix E along with the detailed solution procedure, allows to decouple the dependence of *α*_*c*_ on *γ* and *L*, with the former given by the spatial variance given by Eq. 29 and the latter by the solution of Eq. 26 and 28 in the *γ* = 0 case. We therefore have:

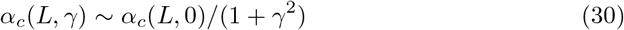

The storage capacity is plotted in Fig. 8 as a function of *γ* and *L*.

**Figure 8:**
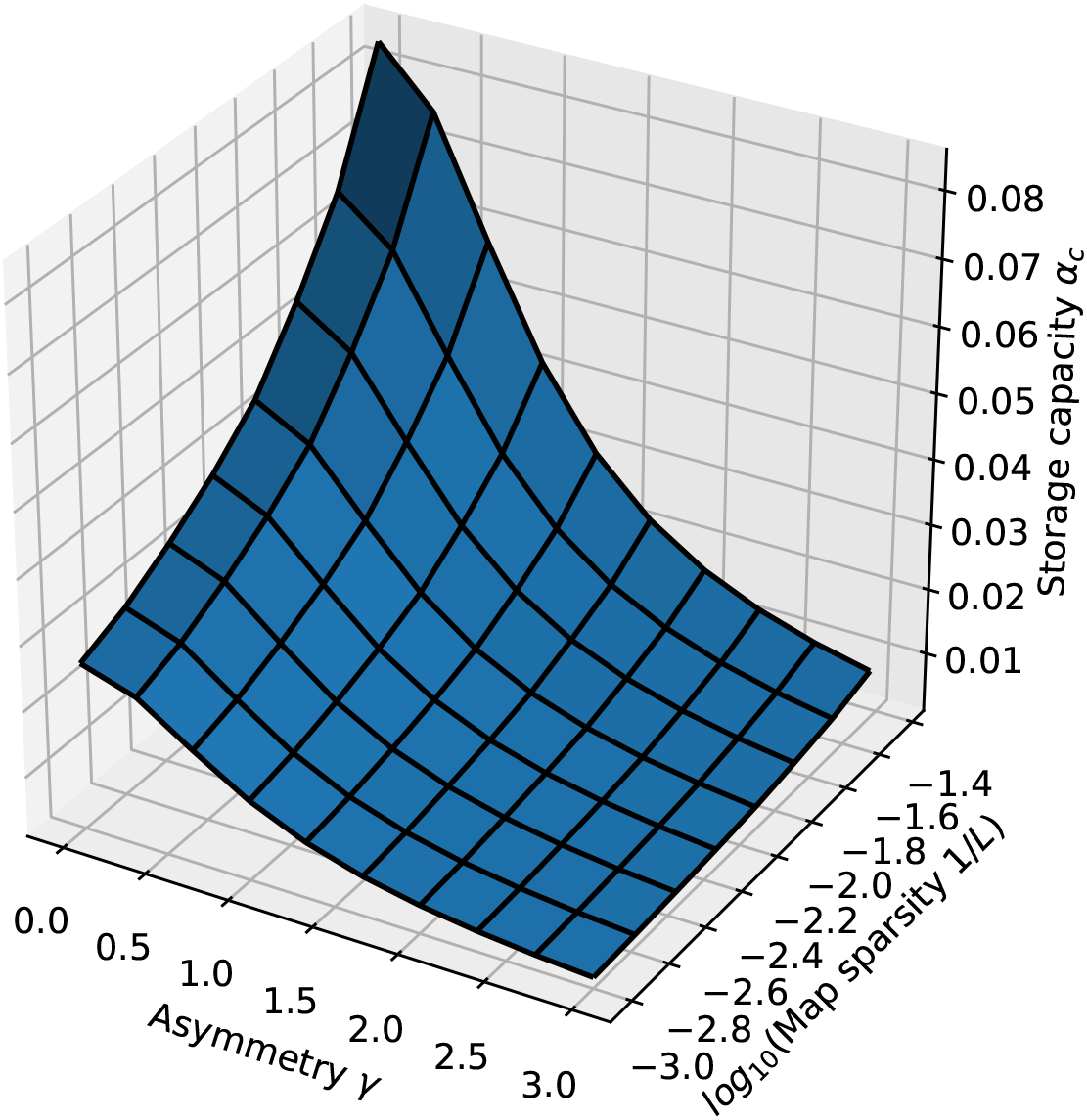
Storage capacity of a diluted network. Dependence of the storage capacity on *γ* and 1*/L* (represented as *log*_10_(1*/L*)).

For sparse maps and small values of the asymmetry, the capacity scales as

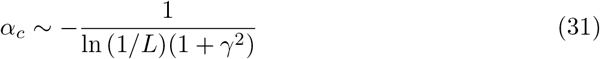

The scaling with 1*/L* is the same found by Battaglia & Treves [30] in the analysis of the symmetric case, as expected: for *γ* = 0 the two models are equivalent.

The presence of asymmetry decreases the capacity, but does not have a catastrophic effect: the decrease is continuous and scales with a power of *γ*. There is therefore a wide range of values of asymmetry and map sparsity in which a large number of dynamic manifolds can be stored and retrieved.

### Numerical estimation of the capacity for a fully connected network

To estimate the storage capacity for a fully connected network, we proceed with numerical simulations. For a network of fixed size *N*, and for given *γ, L* and number of maps *p*, we run a number of simulations *N*_*S*_, letting the network evolve from a random initial configuration. We consider a simulation to have performed a successful retrieval if the global overlap

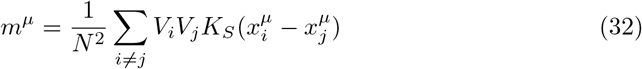

that quantifies the coherence of the activity with map *µ*, is large for one map *µ*\* (at least 95% of the overlap value obtained in the case of a single map) and low in all others maps *µ* ≠ *µ* *. We then define the retrieval probability as *p*_*r*_ = *N*_*R*_*/N*_*S*_, where *N*_*R*_ is the number of observed retrievals.

We repeat the process varying the storage load, i.e. the number of stored manifolds *p*. As *p* is increased, the system reaches a transition point, at which the retrieval probability rapidly goes to zero. This transition is illustrated, for various values of *γ*, in Fig. 9.

**Figure 9:**
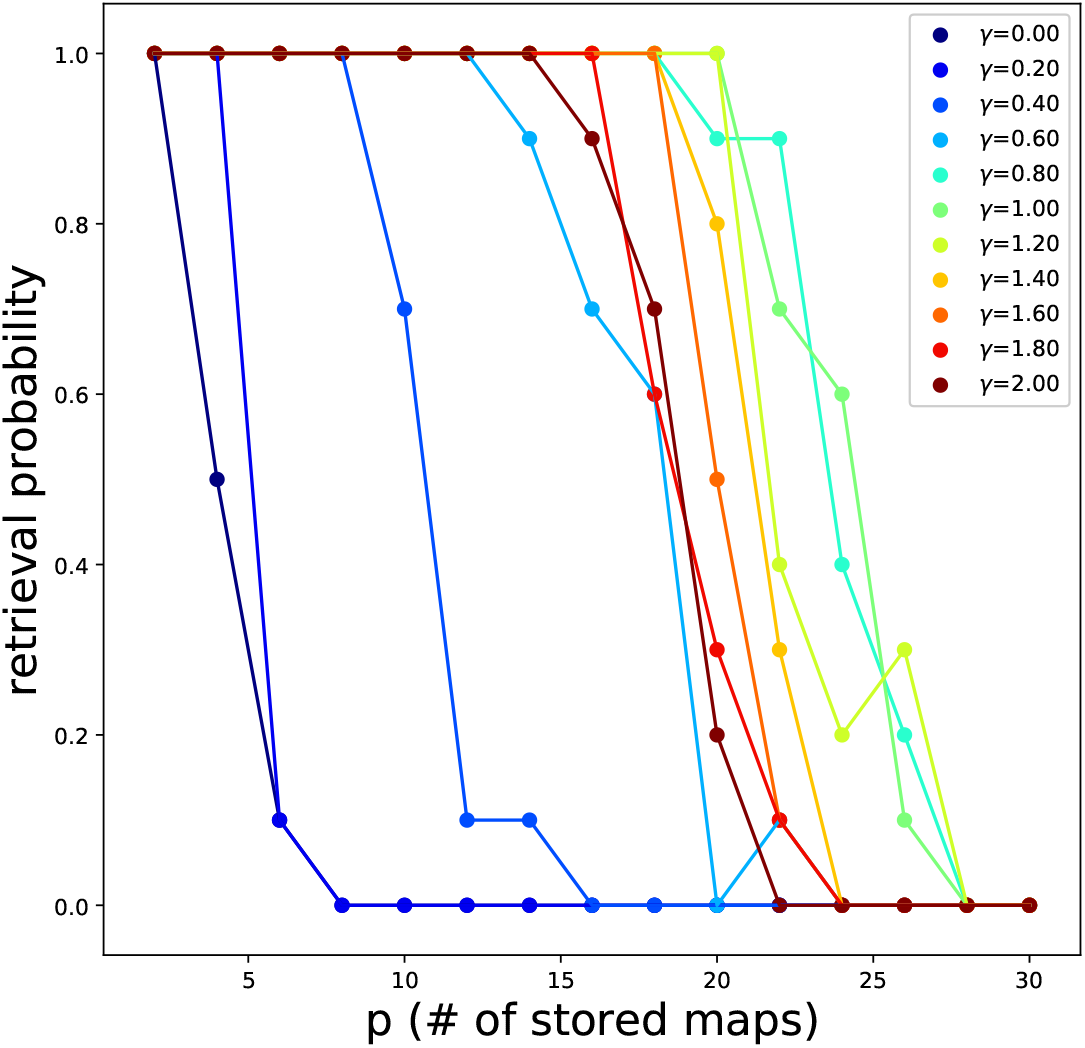
Non monotonic dependence of the capacity on *γ*. Retrieval / no retrieval phase transition for different values of *γ*, obtained from simulations with *N* = 1000, *N*_*S*_ = 10 and *L* = 10, for 1D manifolds. The non-monotonic dependence of the capacity from *γ* can be appreciated here: the transition point moves towards the right with increasing *γ* up to *γ* ∼ 1, then back to the left.

The number of maps *p*_*c*_ at which the probability reaches zero defines the storage capacity *α*_*c*_(*γ, L*) = *p*_*c*_(*γ, L*)*/N*. Repeating this procedure for a range of values of *γ* and *L*, we obtain the plots shown in Fig. 10, for networks encoding one dimensional and two dimensional dynamical memories.

**Figure 10:**
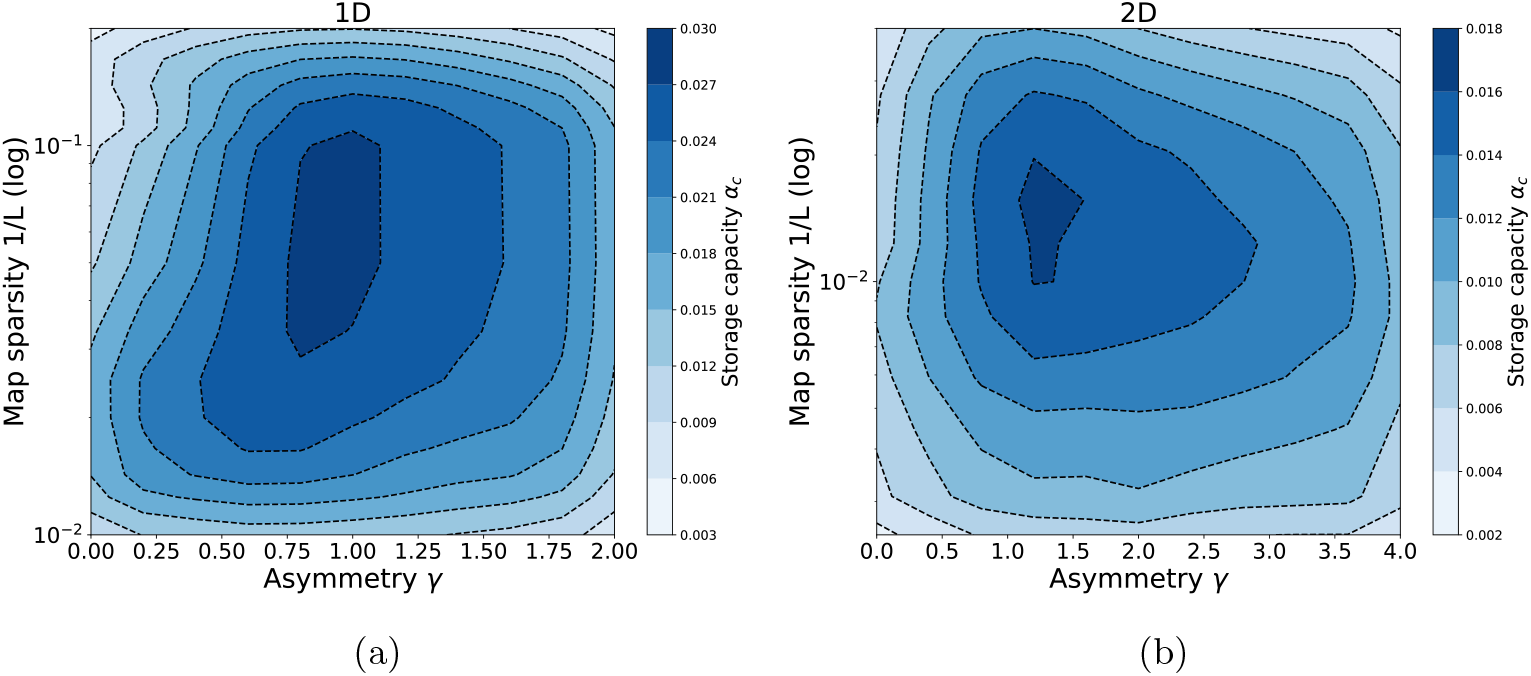
Storage capacity of a fully connected network. Storage capacity as a function of map sparsity 1*/L* and asymmetry strength *γ*, for **(a)** one dimensional dynamic continuous attractors, **(b)** two dimensional dynamic continuous attractors.

The first thing that can be noticed is that, also in the fully connected case, the network can store a large number of maps, for a wide range of *γ* and *L*. A network with size in the order of ten thousand neurons could store from tens up to hundreds of dynamical memories.

The capacity for one dimensional attractors is higher than the one for their two dimensional counterparts. This is in line with what was found for symmetric networks [30].

Finally, we see that the peak of the capacity is found not only for intermediate values of map sparsity – again in line with what is known from the symmetric case – but also for intermediate values of the coefficient *γ*. This shows that moderate values of asymmetry can be beneficial for the storage of multiple continuous attractors, a non-trivial phenomenon that may be crucial for the memory capacity of biological networks. In particular this suggests that the natural tendency of the neural activity to show a rich spontaneous dynamics not only does not hinder the possibility for multiple memories to coexist in the same population, but can be a crucial ingredient for the correct functioning of memory mechanisms.

## Discussion

The results presented show how a continuous attractor neural network with memory-dependent asymmetric components in the connectivity can function as a dynamic memory. Our model is simple enough to be treated analytically, robustly produces dynamic retrieval for a large range of the relevant parameters and shows a storage capacity that is comparable to – and in some cases higher than – the capacity for static continuous attractors.

The analytical solution of the single attractor case shows that the interaction between the strength of the asymmetry and the velocity of the shift can be modulated by global features of the network activity such as its sparsity. This makes the network able to retrieve at different velocities in different regimes, without necessarily requiring short term synaptic modifications. The insensitivity of the general features of the dynamics to the fine details of the shape of the interactions suggests that this mechanism could robustly emerge from learning or self organization processes in the presence of noise. Multiple manifolds can be stored simultaneously in the same neural populations and retrieved at different speeds, and can interact with each other, producing a sequential retrieval of many continuous items. The analysis of the storage capacity shows that the asymmetry does not heavily impair memory performance, and that, in densely connected networks, out of equilibrium effects can be beneficial for memory.

The storage capacity of out of equilibrium continuous attractors has been calculated, in a different scenario, by Zhong et al. [55]. The authors considered the case of an external signal driving the activity bump along the attractor, in a network of binary neurons, and proceeded to calculate the storage capacity with several assumptions that allowed to model the interference of multiple maps as thermal noise. Interestingly, their main result is broadly compatible with what we show here: in the highly diluted regime the velocity of the external signal has a mild – detrimental – effect on the capacity. This hints that out of equilibrium effects could show some form of universality across different network models and implementations of the shift mechanism. Moreover, a high capacity for dynamical sequences has shown to be achievable also in the case of discrete items [56]. Together these results suggest that the introduction of a temporal structure is compatible with the functioning of autoassociative memory in recurrent networks, and open the way to the use of attractor models for the quantitative analysis of complex memory phenomena, such as hippocampal replay and memory schemata.

The model we propose suggests that the tendency of the activity to move in the neural population is a natural feature of networks with asymmetric connectivity, when the asymmetry is organized along a direction in a low dimensional manifold, and that static memories could be the exception rather than the rule. Indeed, Mehta et al. [57] have shown that place fields can become *more* asymmetric in the course of spatial learning, demonstrating that the idea that symmetry emerges from an averaging of trajectory-dependent effects [58] does not always hold true. The model we presented can be useful for the quantification of the effects that symmetry and asymmetry in the interactions have on the acquisition, retention and retrieval of memory.

In most of the two- and three-dimensional cases analysed here the asymmetry is constant along a single direction in each attractor. This can describe the situation in which the temporal evolution of the memory is structured along a certain dimension, and free to diffuse, without energy costs, in the remaining ones. The description of several one-dimensional trajectories, embedded in a two dimensional or three dimensional space requires a position-dependent asymmetric component. A systematic analysis of this situation is left for future analysis. However, the simple case of two intersecting trajectories embedded in a 2D map, analyzed here, provides a proof of concept that several intersecting trajectories can be correctly retrieved, provided that the activity bump is sufficiently elongated in the direction of the trajectory. A progressive elongation of the place fields in the running direction was observed in rats running on a linear track [59], and our analysis predicts that an analogous effect would be observed also in open-field environments, when restricting the analysis to trajectories in the same running direction.

The dynamical retrieval of the model generalizes, in the framework of attractor networks, the idea of cognitive maps, incorporating a temporal organization in the low-dimensional manifold encoding the structure of the memory. This feature is reminiscent of the idea of memory schemata – constructs that can guide and constrain our mental activity when we reminisce about the past, imagine future or fictional scenarios or let our minds free to wander [60]. The use of the present model to describe memory schemata will require further steps, such as an account of the interaction between hippocampus and neocortex, and a mechanism for the transition between different dynamical memories. Nevertheless, the idea of dynamic retrieval of a continuous manifold and the integration of the model presented here with effective models of cortical memory networks [61] open promising perspectives.

Finally, the full analytical description of a densely connected, asymmetric attractor network is a challenge that remains open, and can yield valuable insights on the workings of the neural circuits underlying memory.

## Acknowledgments

Work supported by the Human Frontier Science Program RGP0057/2016 collaboration. We are grateful for inspiring exchanges with Remi Monasson and others in the collaboration, and thank Silvia Girardi for her help with figure 1.

## Appendix

### A Numerical simulations

Numerical simulations are performed with python code, available at: https://github.com/davidespalla/Dynamic-Continuous-Attractors

In the single map case, to each of the *N* units (*N* = 1000 in 1D, *N* = 1600 in 2D) is assigned a preferential firing location *x*_*i*_ on a regular grid spanning the environment with linear dimension *L*. From this preferred firing locations the interaction matrix *J*_*ij*_ is constructed, with the formula:

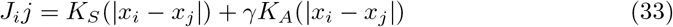

The precise shape of the symmetric and anti-symmetric parts of the kernel are chosen differently in different simulations, according to the feature the analysis focused on, as specified in the main text. Once the network is assembled, the dynamics is initialized either with a random assignment of activity values to each unit in the range [0, 1], or with a gaussian bump centered in the middle of the environment (note that, due to the periodic boundary conditions and the translational invariance of the connectivity, the choice of the starting point does not influence the outcome). The dynamics is then evolved in discrete time steps, with the iteration of the following operations:

- Calculation of the local fields *h*_i_(*t*) = Σ_*i*_ *J*_ij_ *V*_j_ (*t* − 1)
- Calculation of the activity values *V*_*i*_(*t*) = *g*(*h*_*i*_(*t*) − *h*_0_)*θ*(*h*_*i*_(*t*) − *h*_0_)
- Dynamic adjustment of the threshold *h*_0_ such that only the *f N* most active neurons remain active: *h*_0_ = *V*_*i*(*f*)_, where 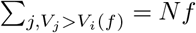
- Recalculation of the activity *V*_*i*_(*t*) with the adjusted threshold
- Dynamic adjustment of the gain *g* such that the mean activity ⟨{*V*_*i*_(*t*)} ⟩ is fixed to 1: *g* = *g/*⟨{*V*_*i*_(*t*)} ⟩
- Recalculation of the activity *V*_*i*_(*t*) with the adjusted gain

The adjustment of the parameters of the transfer function is enforced to constrain the network to operate at fixed sparsity *f* and fixed mean, set to one without loss of generality. The dynamics is iterated for a given number of steps (usually 200), large enough to assure the convergence to the attractive manifold (reached usually in *<* 5 steps) and the observation of the dynamical evolution on the manifold.

In the case of multiple maps, the implemented dynamical evolution is the same, but the interaction matrix is constructed with multiple assignments of the preferred firing locations *x*_*i*_, one for each of the *p* stored maps:

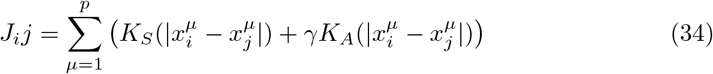

The multiple assignments of the preferred firing locations are performed by a random shuffling of the labels of the units before the assignment to the position on the regular grid spanning each map.

### B Analytical solution of the single map model in one dimension

To solve the integral equation 9 in the case of a 1D manifold and the exponential kernel

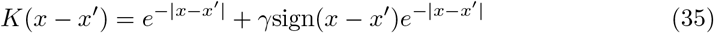

we start by rewriting it as

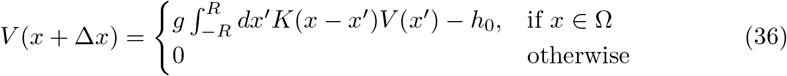

where [−*R, R*], *R >* 0 is a compact domain for which there exist a solution of Eq. 9 that is zero at the boundary. This allows to exploit the fact that our threshold-linear system is, indeed, linear in the region in which *V* (*x*) *>* 0.

We then differentiate twice to obtain the differential equation

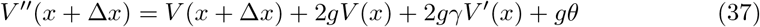

This is a second order linear ODE, with constant coefficients. The presence of the shift term Δ*x* inside the unknown function makes the equation non-trivial to solve. To solve it, we proceed in the following way: first, we look for a particular solution, that is easily found in the constant function

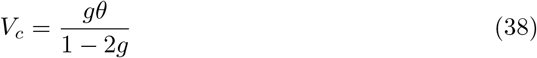

Then, we consider the associated homogeneous equation, and look for a solution in the form *V* (*x*) = *e*^*kx*^, where *k* is a solution of the characteristic equation *C*(*k*) = 0, with

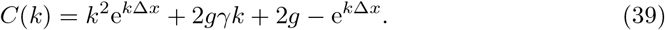

This trascendental equation has to be solved graphically in the complex domain, as shown in Fig. 11.

For each value of *γ* and Δ*x*, the equation shows a pair of complex conjugate solutions

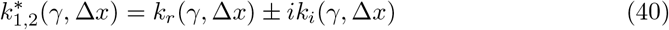

**Figure 11:**
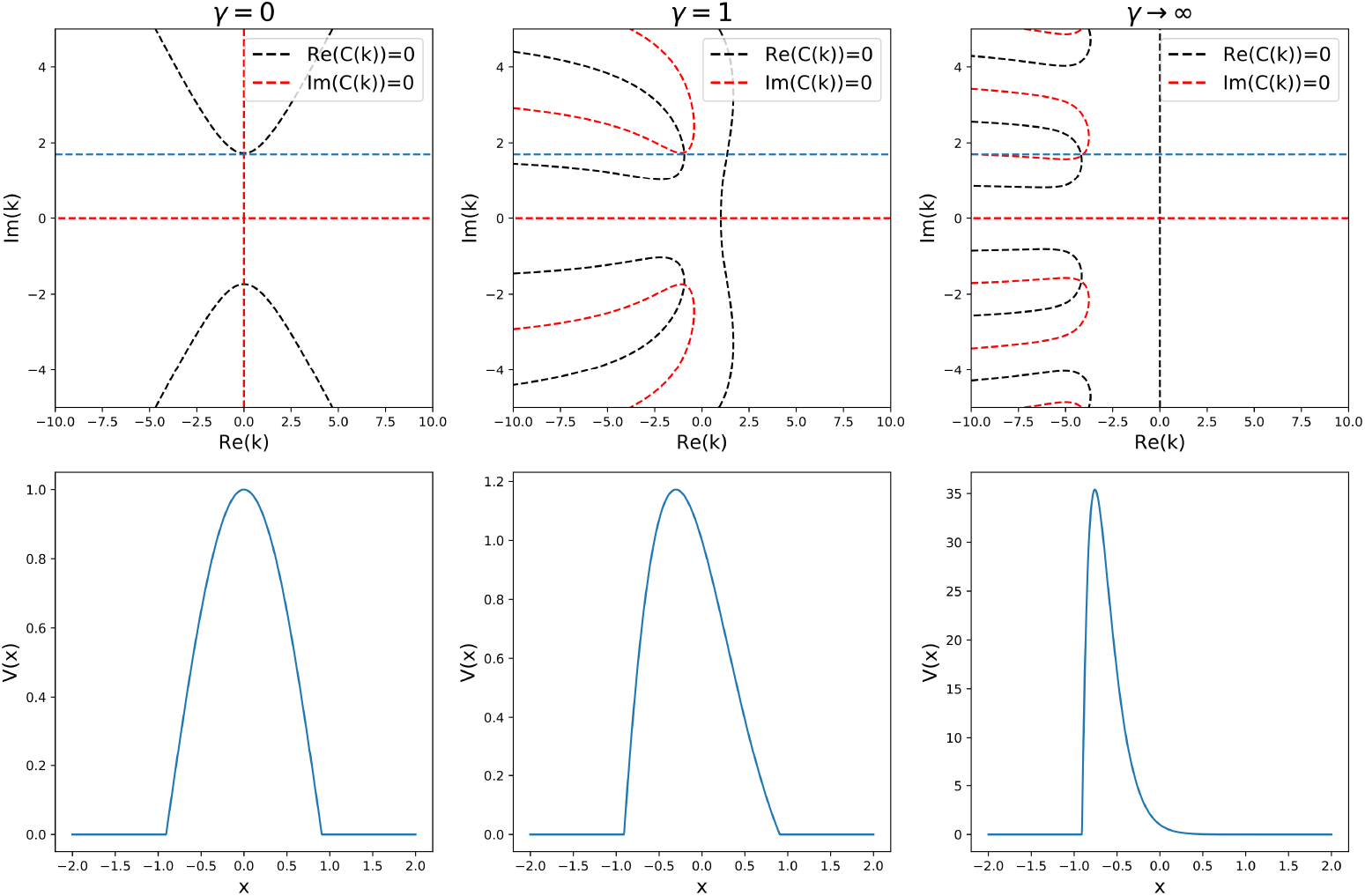
Analytic solution of equation (37).The top row shows the graphical procedure to find the complex zeros of the characteristic *C*(*k*) given in (39), for three different values of *γ*. Black and red lines show the zeros of the real and imaginary part of *C*(*k*), respectively. Their intersections are the complex solutions to *C*(*k*) = 0. The blue line represents the sparsity constraint *k*_*i*_ = *π/*2*R*. The bottom row shows the corresponding solution shapes.

The general solution of the equation will therefore have the form

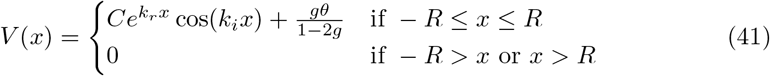

From Eq.41 we can see that the absolute value of *k*_*i*_ is related to the width of the bump, and therefore to the sparsity of the solution, by the relation

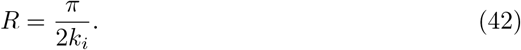

*R* does in turn depend only on the free parameter *g*, through the relation that can be derived in the symmetric case (*γ* = 0, Δ*x* = 0)

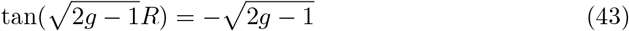

We can look for a solution with given sparsity *f* = 2*R/L* (where *L* is the length of the manifold) by setting *R* = *f L/*2. Requiring the the sparsity to be fixed (i.e. setting *R* constant) while varying *γ* constrains the zeros of 39 to lie in the subspace *k*_*i*_ = *π/*2*R*. This imposes a relation between *γ* and both the speed *s* = Δ*x/τ* (related to the speed of the shift) and *k*_*r*_ (related to the asymmetry of the shape of the solution). Varying *R* we can study the dependence of the speed on both *γ* and *f*.

### C Storing multiple manifolds with different retrieval speeds

To investigate the possibility to store and dynamically retrieve manifolds at different speeds, we have performed numerical simulations of a network with recurrent connectivity given by the formula

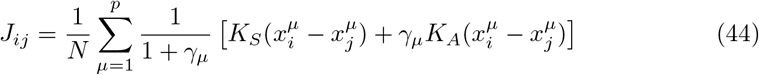

Each manifold *µ* is encoded with a different asymmetry strength *γ*_*µ*_. The normalization factor 1*/*(1 + *γ*_*µ*_) is added to ensure that each manifold contributes equally to the synaptic efficacies, and does not affect the ratio of strenghts of the symmetric and asymmetric components.

### D Linking multiple manifolds together

In order to model the interaction between different stored manifolds, we add to the connectivity matrix an “heteroassociative” term, whose strength 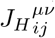 is proportional to the distance between the preferred firing location 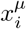 of neuron *i* in manifold *µ* and the one of neuron *j* in manifold *ν*, shifted by the length *L* of the first manifold, which we denote by 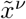. This shift enforces the fact that the interaction happens between the end of the first manifold and the beginning of the second. Then, the connectivity matrix will be given by

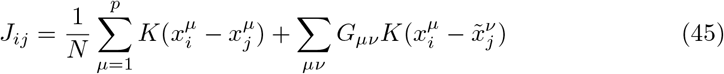

With

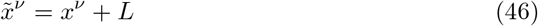

and *G*_*µν*_ = 1 if there is a transition between *µ* and *ν*, and zero otherwise.

### E Analytical calculation of *α*_*c*_ in the highly diluted limit

To calculate the maximum capacity, we first need to solve Eq. 26 numerically for *v*(*x*), for given *g* and *w*. The procedure illustrated here focuses, for the sake of analytical simplicity, on the case of a one-dimensional, exponential kernel

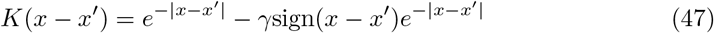

We start from equation 26:

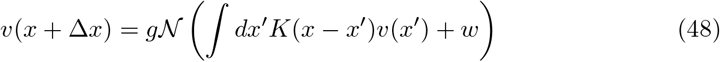

First, following [30] we rewrite it with the transformation

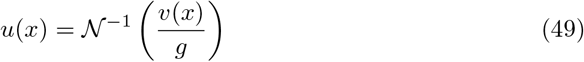

obtaining

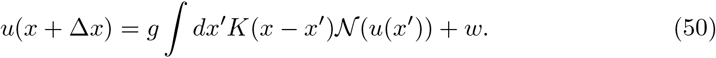

We then transform this integral equation in a differential one, by differentiating twice. We obtain

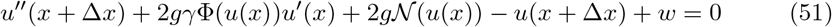

where we have used the fact that 𝒩′ (*x*) = Φ(*x*). Eq.51 is a second order, nonlinear delayed differential equation. To solve it, it is not sufficient to impose an initial condition on a single point for the solution and the first derivative (i.e. something like 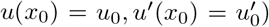): we have to specify the value of the function and its derivative in an interval [*x*_0_, *x*_0_ + Δ*x*].

To do so, we reason that, if we want a bump solution, *u*(*x*) has to be finite for *x*→ ± ∞ and cannot diverge. We then require the function to be constant (*u*(*x*) = *u*_0_, *u*′ (*x*) = 0) before a certain value *x*_0_, whose value can be set arbitrarily without loss of generality.

The value *u*_0_, at *γ* = 0 and Δ*x* = 0 determines the shape of *u*(*x*), as shown by the numerical solution presented in Fig. 12. For *u*_0_ *< u*\* the solution will diverge at *x*→ ∞, while for *u*_0_ *> u*\* it will oscillate. We are then left with a single value *u*_0_(*g, w*) = *u*\* (*g, w*) for which the solution has the required form.

**Figure 12:**
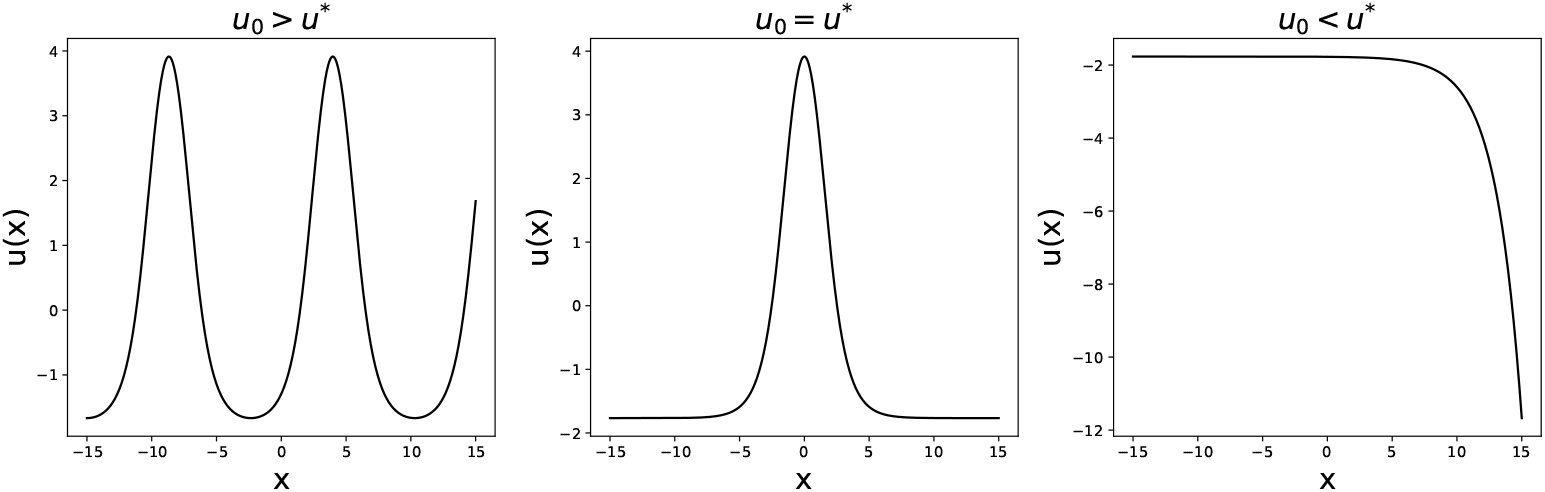
Solutions to Eq.51 for *g* = 1, *w* = −1.8, *γ* = 0, Δ*x* = 0.

Then, keeping *u*_0_ fixed, we can repeat a similar procedure to find Δ*x* for different values of *γ*. Also in this case, the solution either diverges or oscillates, apart from a single value Δ*x*\*, for which the solution has the desired shape (see Fig.13). This eliminates the arbitrariness in the choice of Δ*x* since it imposes, for given *g* and *w*, a relation Δ*x* = Δ*x*\* (*γ*).

**Figure 13:**
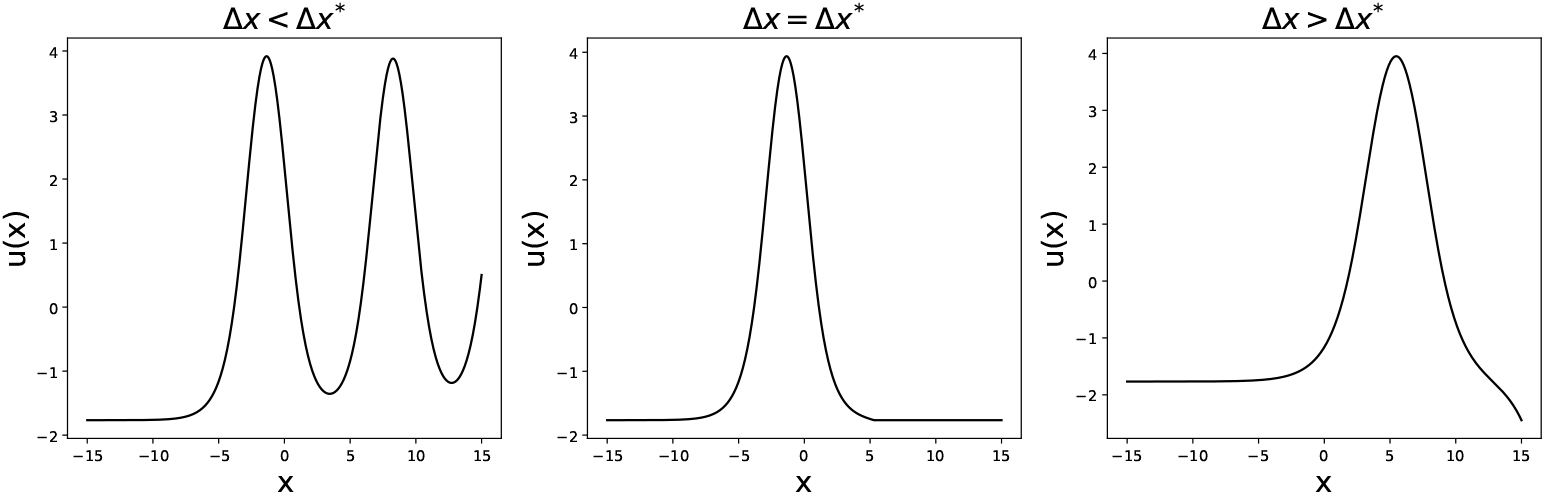
Solutions to Eq.51 for *g* = 1, *w* = −1.8, *γ* = 0.2.

We can then find the shape of the bump *u*(*x*) for given values of *g*, +*w* and *γ*, from which we can obtain the profile *v*(*x*) = *g*𝒩 (*u*(*x*)) that we need for the calculation of the storage capacity. Some examples of the obtained profiles, for different values of *γ*, are shown in Fig. 14.

**Figure 14:**
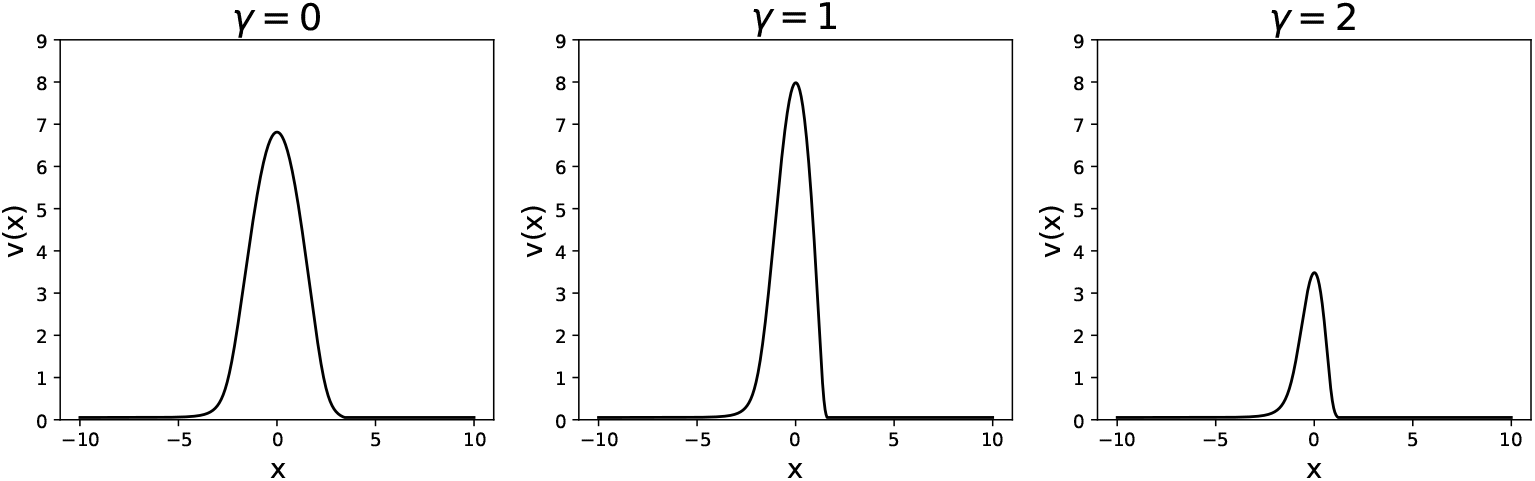
Activity profile *v*(*x*), obtained for the same *g* = 0.7 and *w* = − 1.3, at different values of *γ*.

Plugging the obtained form of *v*(*x*) into Eq. 28, we can calculate the capacity. The dependence of the capacity on *γ* is shown, for *L* = 60, in Fig. 15.

**Figure 15:**
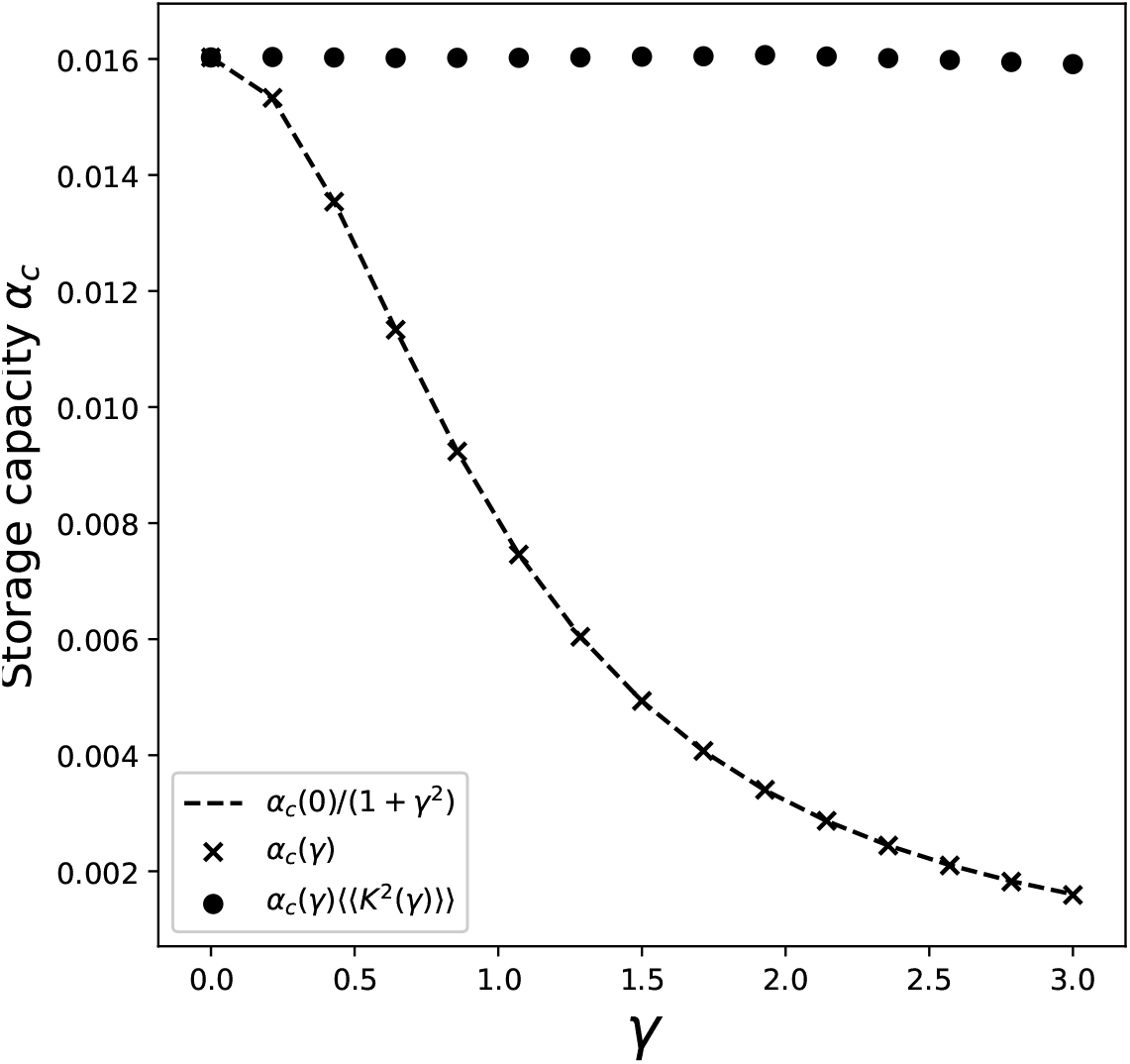
Dependence of the storage capacity on *γ*, for *L* = 60. The crosses show the full solution of Eq. 26 and 28. The dashed line is obtained by taking the value of the capacity *α*(0) obtained with full solution at *γ* = 0, and multiplying it by the scaling of the kernel variance (1 + *γ*^2^). Full dots show the value of capacity obtained with the full solution and the contribution of the kernel variance factored out.

We can see from the full dots in the figure that the contribution of the integral in Eq.28 is remarkably constant in *γ*. This is due to the fact that the distortions of the bump shape induced by the presence of the asymmetry have a negligible effect on the average square activity *y*, whose value is dominated by the dependence on *γ* of the spatial variance of the kernel (Eq.18).

This allows us to approximate the value of the integral in Eq. 28 with its value in the *γ* = 0 case. We can then calculate the capacity as a function of *γ* and *L* by solving the symmetric case for different *L*s, and then incorporating the dependence on *γ* given by the kernel variance:

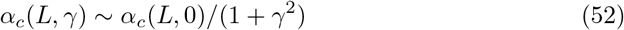

This approximation yields the results reported in the main text and in Fig. 8

